# Opioid Modulation of Differential Gene Expression and Neuronal Differentiation in the Ventricular-Subventricular Zone of Adult Male Zebra Finches

**DOI:** 10.1101/2024.05.20.595080

**Authors:** Uzma Din, Sourav Banerjee, Soumya Iyengar

## Abstract

The endogenous opioid system modulates diverse functions including pain, physiological functions and motivation. We had earlier demonstrated an increase in cell proliferation in the neurogenic niche of adult male zebra finches, following administration of the general opioid antagonist, naloxone. Since the mechanisms underlying opioid modulation of neurogenesis are poorly understood, we have investigated the molecular mechanisms underlying cell proliferation differentiation induced by the opioid system. Systemic naloxone or vehicle administration in adult male birds was followed by a whole transcriptome microarray assay of the ventricular-subventricular zone (V-SVZ). The analysis revealed 26 differentially expressed transcripts (expression fold change ≥ 1.5), encoding genes important for neuronal functions. The transcriptomics analysis using microarray identified a significant increase in the expression of precursor miRNA *tgu-mir-124-201* transcript following inhibition of opioid receptors by naloxone. Our results suggested that the administration of naloxone triggers an increase in the expression of the pro-differentiating microRNA, miR-124, in the brains of adult zebra finches. Using quantitative real-time PCR, we found that the expression of miR-124-3p was higher in the V-SVZ of naloxone-treated versus control birds. Furthermore, the density of neuroblasts was higher in the ventral V-SVZ adjacent to the striatal nucleus Area X, but not in the V-SVZ above the pallial nucleus, HVC, both of which are involved in singing. Since miR-124 targets signalling pathways important for neuronal proliferation and differentiation, our findings suggest that microRNA-124 may be involved in the increase in neurogenesis in adult songbirds induced by altering opioid modulation.

## 2 Introduction

The two main niches that serve as the epicentres for adult neurogenesis in the vertebrate brain are the ventricular-subventricular zone (V-SVZ), lining the lateral ventricles in the telencephalon [1] and the subgranular zone (SGZ) within the dentate gyrus of the hippocampus [2]. Whereas the rate of generation of new neurons in the V-SVZ varies among vertebrate species from fishes to primates [3], [4], the magnitude of neurogenesis in the V-SVZ of adult songbirds such as zebra finches (*Taenopygia guttata*) is significantly higher than that of other species. In these birds, nascent neurons migrate and incorporate into various parts of the telencephalon, including some of the song control nuclei [5], [6], [7]. Earlier studies from our lab [8], [9] demonstrated that opioid receptors (ORs) were expressed by the V-SVZ of adult zebra finches and blocking these receptors with systemic injections of the general opioid agonist naloxone led to an increase in cell proliferation *in vivo*. Similar results were observed when primary cultures of the adult zebra finch V-SVZ were treated with naloxone [10]. The aim of the present study was to identify the genes involved in the regulation of neurogenesis in the V-SVZ of adult male zebra finches following systemic injections of naloxone. To elucidate differential gene expression patterns induced by naloxone administration, a whole transcriptome microarray analysis was performed of the V-SVZ of adult male zebra finches. Besides transcripts encoding genes important for neuronal function and cell proliferation, we found a significant increase in the upregulation of a transcript for the precursor miRNA tgu-mir-124-201. These findings are interesting since miR-124 inhibits cell proliferation and promotes differentiation into a neuronal fate via SRY-box transcription factor Sox9 [11], [12]. Furthermore, miR-124 represses extracellular matrix proteins LAMC1 and ITGB1, leading to the differentiation of neuronal precursors into neurons [13]. Our findings therefore suggest that naloxone triggers an increase in the expression of the pro-differentiating microRNA, miR-124, in the neurogenic niche in adult zebra finches. This molecular alteration induced by opioid intervention may direct the newly proliferated cells towards differentiation into neuronal phenotypes.

## 3 Methods

### 3.1 Animals and injections

All experimental procedures were performed in agreement with the recommendations of the Institutional Animal Ethics Committee of the National Brain Research Centre, Manesar (NBRC) in accordance with the guidelines of the Committee for Control and Supervision of Experiments on Animals (CCSEA), India, which comply with international guidelines for animal welfare. A total of 32 adult male zebra finches (*Taeniopygia guttata*; >120 days post hatch) were maintained in aviaries exposed to 12L:12D cycles of light and dark and provided ad libitum access to feed (millets, eggs, greens) and water. Nine birds received intramuscular injections of 2.5 mg/kg body weight of the opioid antagonist naloxone (Naloxone hydrochloride dihydrate, Sigma, cat no. N7758) for four consecutive days, whereas controls (n = 9) were injected with 0.9% saline (vehicle) during the same period. For histology and immunolabelling experiments, another set of 3-4 birds were injected with naloxone or saline as described above, followed by an injection of 100 mg/kg body weight BrdU (Bromodeoxyuridine, B9258, Sigma). This set of birds were sacrificed using an overdose of ketamine hydrochloride on the fourth day, two hours after the last injection.

### 3.2 Sample preparation, pre-processing and whole transcriptome microarray assay

For microarray and qRT-PCR experiments, sample preparation and pre-processing steps were performed under sterile, RNAse-free conditions. Birds were overdosed with halothane, after which brains were removed and cut into 1mm thick coronal slices. Using a dissection microscope (Carl Zeiss™ Stemi 2000-C), a region approximately 250µm lateral to the lateral ventricles was dissected from these slices, containing both the ventricular and subventricular zones (cf. Khurshid et al., 2010). For both treated and control groups, V-SVZ from three birds were pooled and total RNA was isolated by the guanidinium thiocyanate-phenol-chloroform or TRIzol method. The extracted RNA was treated with TURBO™ DNase (Invitrogen, cat no. AM2388) to remove any traces of genomic DNA, quantified and checked for RNA integrity (RIN ≥ 7). All procedures involved in the microarray study were performed at Innovative Life Discoveries Private Limited, Manesar, India. For each sample, 500ng of the total RNA was processed and hybridized to a 64-format Affymetrix Genechip Zebra Finch 1.0 ST Array, using standard protocols provided in the WT PLUS reagent kit by Affymetrix®. Staining, washing and scanning of the array were performed in accordance with standard Affymetrix protocols. Raw data files were processed for a background adjustment, quantile normalization and summarisation in the Affymetrix Expression Console. The filtered probes were annotated using the Ensembl based taeGut3.2.4 assembly.

### 3.3 Identification of Differentially Expressed Genes and Hierarchical Cluster Analysis

A moderated t-test was applied to the signal intensities of the filtered probes, for both control (n = 9) and naloxone-treated groups (n = 9; 3 sets of repeats). To consider a gene as differentially expressed and statistically significant, the p value <0.05 and log2 (Fold change) ≥0.59216535 were selected as the cut-off points and represented by Volcano plots. The normalized intensities of the differentially expressed and statistically significant genes with fold change ≥1.5 were input in the Genesis© tool (http://genome.tugraz.at/) and a heat map dendrogram was generated.

### 3.4 Quantitative Real Time Polymerase Chain (qRT-PCR) analysis

For further validation by qRT-PCR, we selected differentially expressed transcripts based on the genes encoding identifiable proteins or miRNAs, particularly those participating in pathways related to cell division and differentiation. Specific Taqman™ microRNA assays (Assay ID: 000239, cat. no. 4427975) and miR-124-5p (Assay ID: 241919_mat, cat. no. - 4440886) and specific primer sequences were used to detect the genes of interest by qRT-PCR (Table-4). The detection of miR-124 was performed adhering to manufacturer’s instructions from Taqman™ (Invitrogen) with the U6 snRNA 3p (Assay ID: 001973, cat. no. 4427975) serving as the reference standard gene. For other genes of interest, the glyceraldehyde 3-phosphate dehydrogenase (GAPDH) gene was used as an endogenous control. The relative amount of each gene of interest was normalized against U6 snRNA or GAPDH, and the fold change (FC) for each gene in naloxone-treated samples compared with vehicle controls was calculated by using the 2−ΔΔCt method.

### 3.5 Histology, immunostaining, imaging and colocalization analysis

Following sacrifice and perfusion of the birds, brains were removed, cryopreserved in 30% sucrose and sectioned coronally into 30 µm thick serial sections using a cryostat (CM 3050 S, Leica). To identify song control nuclei and other anatomical markers, one series of the sections was stained with Thionin (Nissl). For analysing the density of newly proliferated neuroblasts, three sections were selected from each bird, spanning the coronal planes containing the V-SVZ adjacent (i) Area X (n = 4) and (ii) HVC (n = 3; [14]. The sections were rinsed in PBS, subjected to antigen retrieval (Vector Laboratories, H-3300, pH 6) and treated with 2N HCl, to denature DNA. Sections were then rinsed in 0.1M Borate buffer, followed by quenching with 1% hydrogen peroxide and rinsed in 0.1% Triton-X-PBS. Next, sections were blocked in 10% Normal goat serum (NGS, Vector Laboratories, S-1000), 2% Bovine serum albumin (BSA), and 0.1% Triton-X-PBS for 2 hours. The sections were then incubated in a solution containing mouse monoclonal anti-BrdU antibody (1:250, Sigma, Cat.no. B8434) and rabbit polyclonal anti-Doublecortin (DCX) antibody (1:1000, Abcam, ab18273, [15] diluted in 5% NGS, 2% BSA, and 0.1% Triton-X-PBS at 4°C for 36-48 hours. Following PBS washes, the sections were incubated with a cocktail of secondary antibodies [goat anti-mouse Alexa-488 (1:500, Vector Laboratories, A11001) and goat anti-rabbit Alexa-594 (1:1500, Vector Laboratories, A11012)], for 2 hours at room temperature. After rinsing with PBS, sections were mounted on gelatin-coated slides and cover-slipped using Vectashield® Antifade mounting medium with DAPI (Vector Laboratories, H-2000).

For each immunolabelled section, images of BrdU-positive nuclei and DCX-positive neuroblasts within the region extending up to approximately 185µm lateral to the V-SVZ [10], [14] were acquired as a series of z-stacks at 60x, with a step size of 0.5 µm and a resolution of 1024x1024 pixels using a Nikon Confocal microscope (A1 HD25). Newly proliferated groups of BrdU-positive cells were quantified in maximum projection images generated from the z-stacks using a custom-designed cell counting macro in the ImageJ software. A semi-automated object-based colocalization analysis (OBCA) method [16] was used to count BrdU-positive cells co-labelled with DCX. For each bird, the density of newly proliferated cells and neuroblasts was averaged in three sections at the level of Area X and HVC and presented as the density of BrdU-positive cells and BrdU-DCX positive cells per mm^2^ of these regions.

### 3.6 Statistics

For qRT-PCR experiments, Student’s t-test (unpaired, two-tailed) was performed on the relative fold changes in GraphPad Prism (version 10.0.0, GraphPad Software, Boston, Massachusetts USA) to analyse differences in normalized gene expression. The statistical comparisons between the means of the counts from the two groups were made using an unpaired t-test (two-tailed) in GraphPad Prism. The data were represented as mean ± standard error (SEM), and the significance level was set at p < 0.05.

## 4 Results

### 4.1 Differentially Expressed Transcripts

Microarray analysis of total RNA was performed from the V-SVZ of naloxone-treated birds and control birds and the differentially expressed transcripts determined (**Supplementary Tables 1-3**). Based on the threshold of P values < 0.05 and Log2 fold change (FC) ≥ 0.59216535, that is, FC ≥ 1.5, the expression of 16 transcripts was significantly upregulated (**Supplementary Table 2**) whereas that of 10 transcripts was significantly down-regulated in V-SVZ (**Supplementary Table 3**), following naloxone treatment (**Figure 1**). The Ensembl IDs of the transcripts and the genes encoding them along with their expression fold changes are summarized in **Supplementary Tables 2** and **3**.

**Figure 1.**
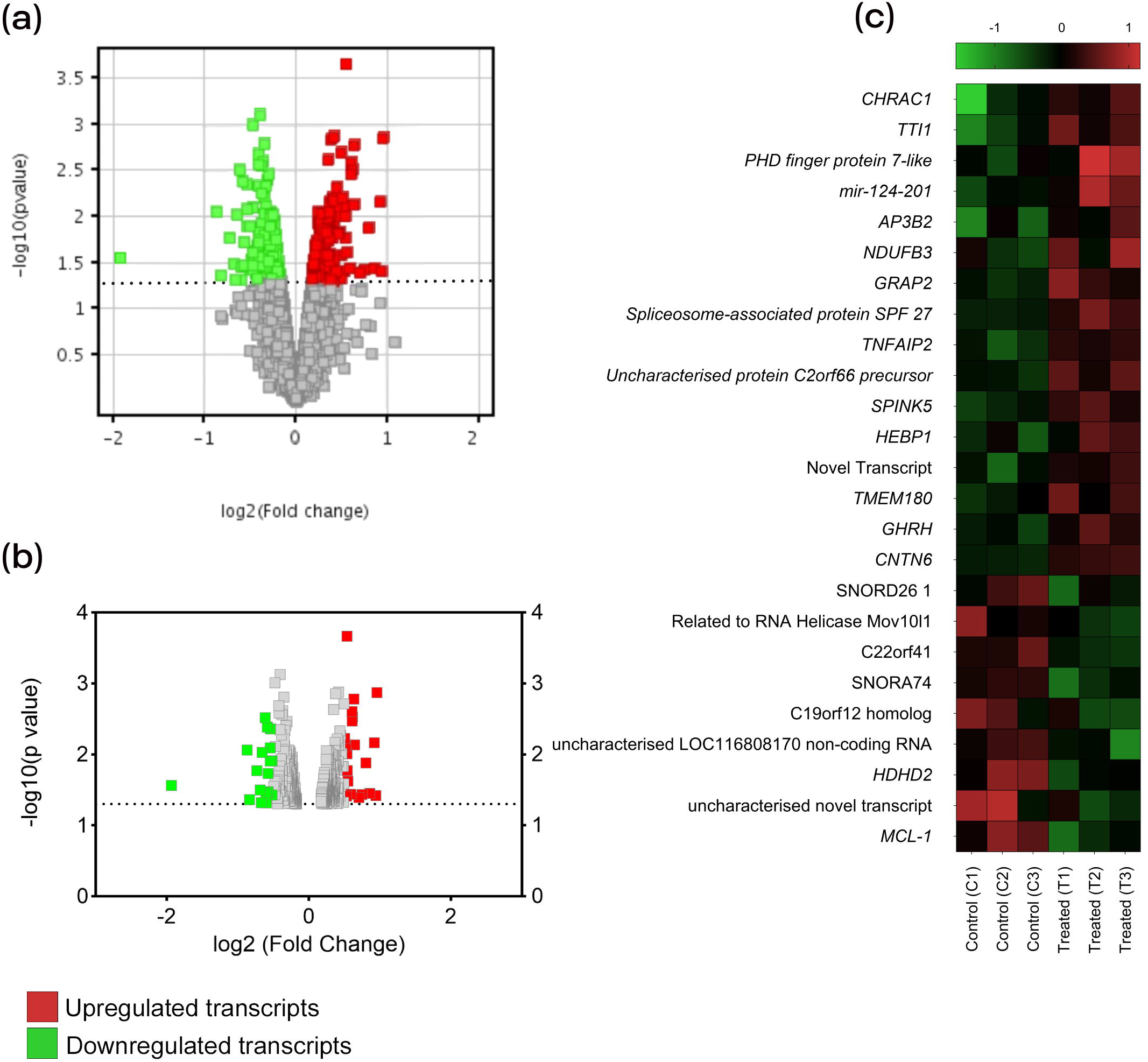
Identification of statistically significant and differentially expressed transcripts. (**a**) Out of 22,220 filtered probe sets, 427 transcripts were revealed to be statistically significant and differentially expressed. Red and green points represent differentially expressed genes (DEGs) with -log10 (P value) ≥ 1.301029, and the grey points (below the dotted line) depict non-significant DEGs. (**b**) After setting the fold change cut-off at |log2 (fold change)| ≥ 0.59216535 or FC ≥ 1.5, 33 transcripts were identified as being the most differentially expressed among the 427 transcripts. Red and green points represent upregulated and downregulated transcripts, respectively. (**c**) Heat map visualization of differentially expressed transcripts with fold change (FC) ≥ 1.5, depicting normalized intensity values. Red squares depict over-expression and green squares represent under-expression of a particular transcript. Columns: Three biological replicates for two experimental conditions; Control (C) and naloxone-treated (T); (n = 9 control and 9 treated birds), Rows: Transcripts.

### 4.2 Blocking Opioid Receptors Leads to an Upregulation of Genes that Influence Neuronal Function in the V-SVZ

The transcripts that are significantly upregulated in the V-SVZ of the birds treated with naloxone are known to be encoded by genes involved in biological functions, including cell-cell signalling, mRNA processing, chromatin remodelling, cell division, cell growth and survival, regulation of apoptosis, protein transport, and exocytosis. For further validation by qRT-PCR, we selected transcripts based on the genes encoding identifiable proteins or miRNAs, particularly those linked to cell division and differentiation pathways (**Supplementary Table 4**). The transcripts upregulated following naloxone administration included TNF alpha induced protein 2 (*TNFAIP2*; FC = 1.53), Adaptor Related Protein Complex 3 Subunit beta 2 (*AP3B2*; FC = 1.7), pre-miRNA-124 (*mir-124-201*; FC = 1.7), TELO2 interacting protein 1 (*TTI1*; FC = 1.9) and Chromatin accessibility complex subunit 1 (*CHRAC1*; FC = 1.9), whereas the Induced myeloid leukemia cell differentiation protein (*MCL-1*; FC = 1.8) was significantly downregulated (**Figure 2a-g**). A quantitative RT-PCR analysis of the upregulated genes, including *miR-124-3p, TELO2, CHRAC-1*, and *AP3B2*, revealed that their expression in fold changes mirrored the results obtained from the microarray analysis. Furthermore, we observed higher levels of *TNFAIP-2* in the V-SVZ of naloxone-treated birds compared to controls, although these differences did not reach statistical significance. Additionally, despite the microarray analysis indicating a significant downregulation of the MCL-1 transcript (FC = 1.8), the qRT-PCR assay demonstrated no changes in its expression levels in the V-SVZ of naloxone-treated birds compared to controls.

**Figure 2.**
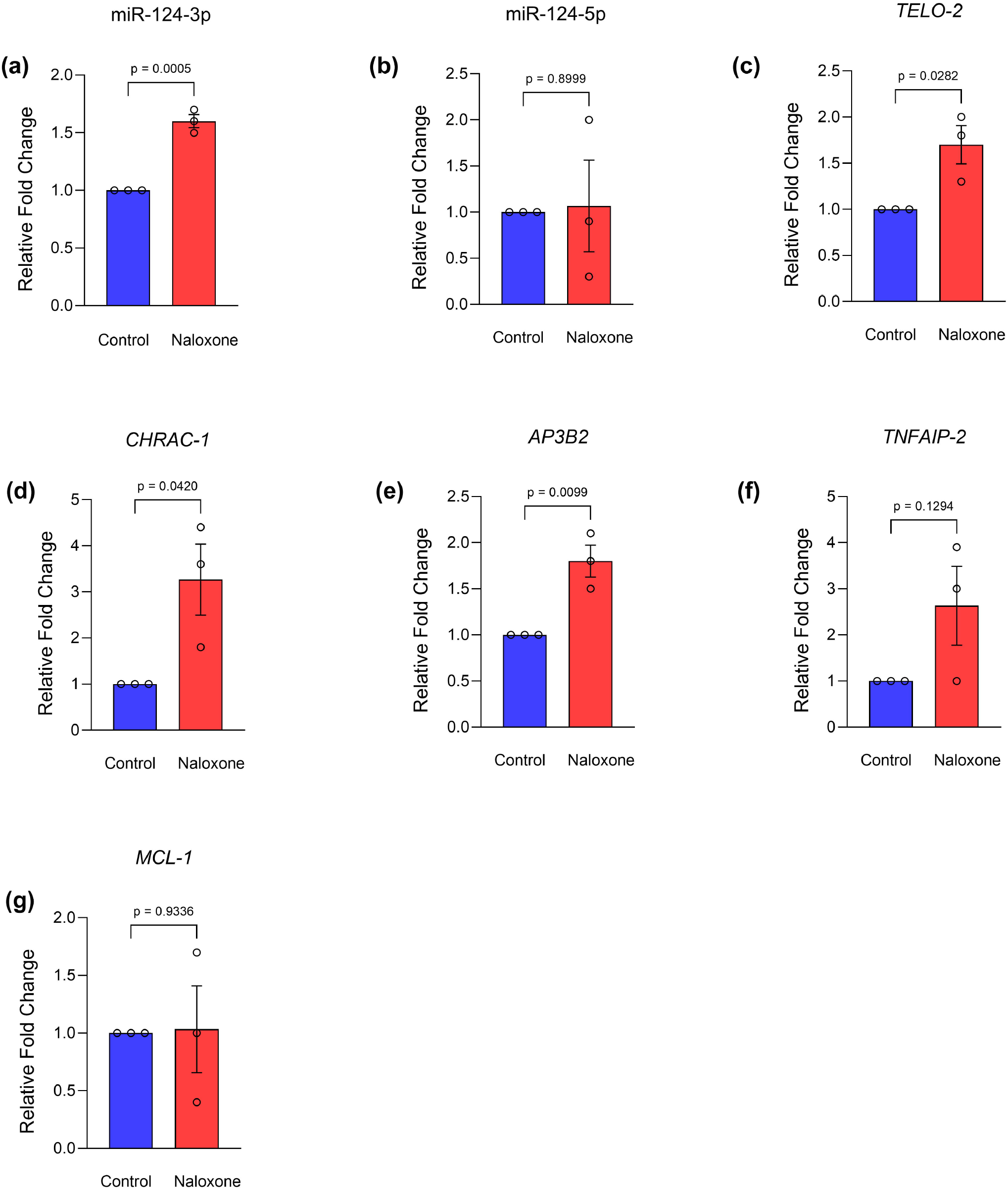
Validation of fold change expression by qRT-PCR. Blocking opioid receptors with naloxone upregulates genes that influence neuronal function and neurogenesis in the V-SVZ. Fold changes in response to naloxone administration for (**a**) miR-124-3p (1.6 ± 0.05); (**b**) miR-124-5p (1.1 ± 0.49); (**c**) Telo-2 Interacting Protein (1.7 ± 0.2); (**d**) Chromatin Accessibility Complex Subunit 1 (3.2 ± 0.76); (**e**) Adaptor Related Protein Complex 3 Subunit beta 2 (1.8 ± 0.17); (**f**) Tumor necrosis factor alpha-inducible protein 2 *(2*.*6* ± 0.85) and (**g**) Induced myeloid leukemia cell differentiation protein (1.03 ± 0.33) are shown versus that in controls, which is assumed to be 1.0. Unpaired t-test, bars represent fold change = means ± SE; *P < 0.05 vs. control, n = 9 controls and 9 treated birds (3 replicates each for control and treated birds)

### 4.3 MiR-124 may be involved in the modulation of naloxone-mediated neurogenesis in the zebra finch V-SVZ

We found a precursor miRNA transcript, that is, tgu-mir-124-201 (miRNA Entry for MI0013699; https://mirbase.org/hairpin/MI0013699), upregulated in naloxone-treated birds versus controls, in the whole transcriptome profile. We decided to validate and quantify mature miRNA-124 levels in the naloxone-treated V-SVZ by qRT-PCR, and compare it with that in controls. Quantitative RT-PCR analysis of both tgu-miR-124-3p and tgu-miR-124-5p revealed that the expression of the 3p strand was significantly upregulated, whereas there was no significant difference in the expression of the miR-124-5p strand in the V-SVZ of naloxone-treated birds compared with that in controls (**Figures 2a** and **2b**), suggesting that miR-124-3p and not miR-124-5p, is the functional and more stable form of microRNA in the V-SVZ after naloxone treatment.

### 4.4 Effects of naloxone on neuronal differentiation in the adult zebra finch V-SVZ

Previous studies have reported that the zones of maximum cell proliferation called ‘hotspots’ are located in the VZ adjoining the song control nuclei Area X and HVC and the anterior commissure in male zebra finches [14], [17]. Our present results demonstrated that the ventral V-SVZ (VVZ) adjacent to Area X and the intermediate V-SVZ (IVZ) dorsal to HVC contained BrdU-labelled cells (green, *arrowheads*) and neuronal precursors co-labelled for BrdU and DCX (yellow, *arrows*, **Figure 3 and 4**, respectively). The density of BrdU-labelled cells and that of BrdU-DCX co-labelled cells was significantly higher in VVZ adjacent to Area X of the naloxone-treated group compared to that in controls (n = 4, p = 0.0026, f = 6). Significant differences were also found in the density of neuronal precursor cells (n = 4, p =0.0463, t = 4.923, df = 6) at this level in naloxone-treated birds. Despite an increase in cell proliferation in the IVZ adjacent HVC, we found that there were no significant differences in the density of BrdU-positive cells (n = 3) or BrdU-DCX co-labelled cells (n = 3) in naloxone-treated versus control birds at this level.

**Figure 3.**
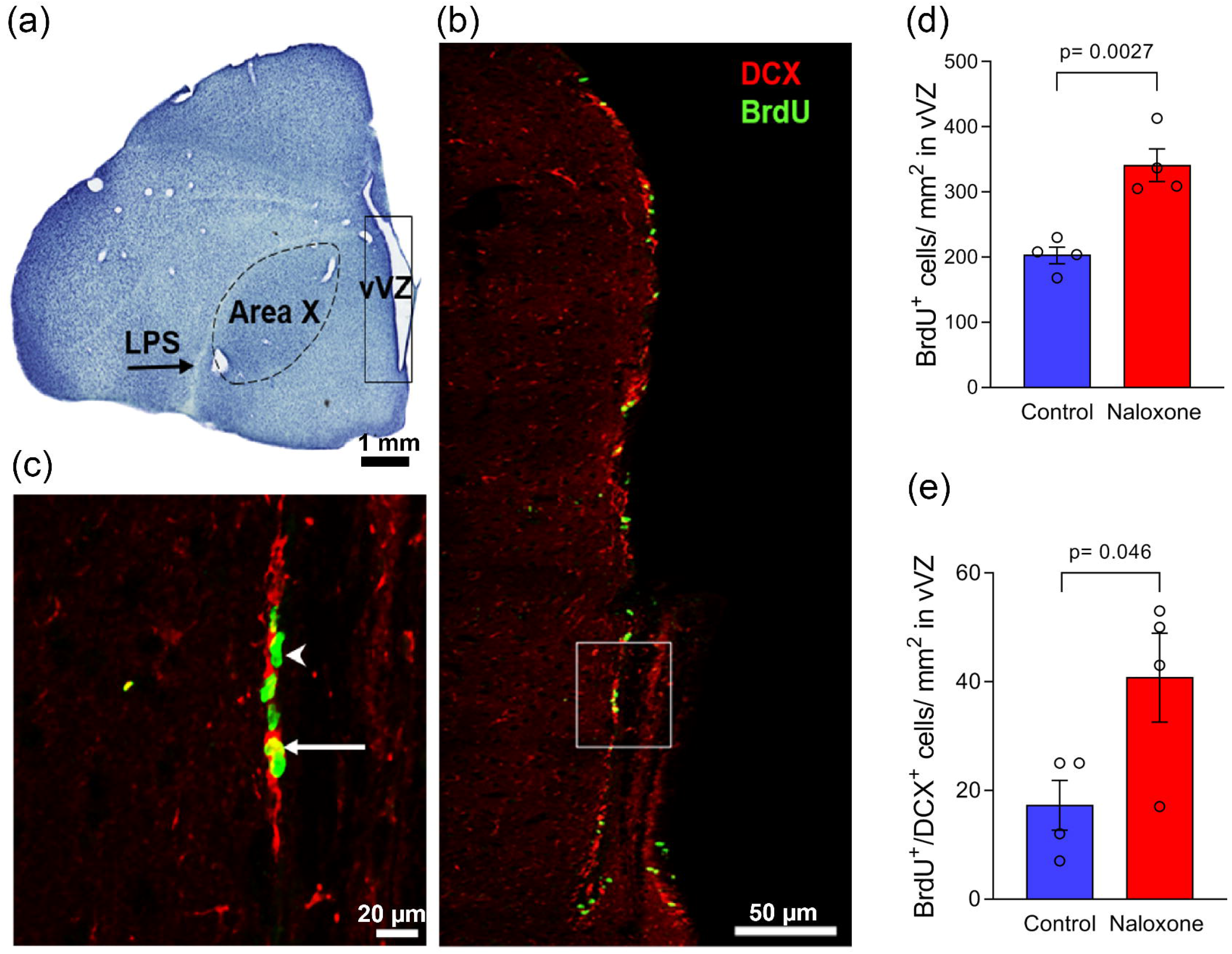
Opioid modulation of proliferation and neuronal differentiation in the VVZ at the level of Area X. **(a)** A Nissl-stained coronal section of the zebra finch brain at the level of Area X demonstrating the ventral part of the ventral ventricular zone (VVZ; black rectangle), from which cells of interest were counted; scale bar = 1mm. **(b)** Representative confocal images at this level, demonstrating DCX-positive neuronal precursors (red) co-labeled with the S-phase marker, BrdU (green); scale bar = 50 µm. **(c)** A high magnification view of the inset (white rectangle in **b**) demonstrates a BrdU-positive cell in the VVZ (*arrowhead*) and DCX-BrdU co-labeled cells (*arrow*); scale bar = 20µm. There were significant increases **(d)** in the density of BrdU+ cells/ mm^2^ (n = 4) and **(e)** co-labeled BrdU/DCX cells/ mm^2^ (n = 4) in the VVZ of naloxone-treated birds versus that in controls (Unpaired t-test, bars represent means ± SE, p<0.05).

**Figure 4.**
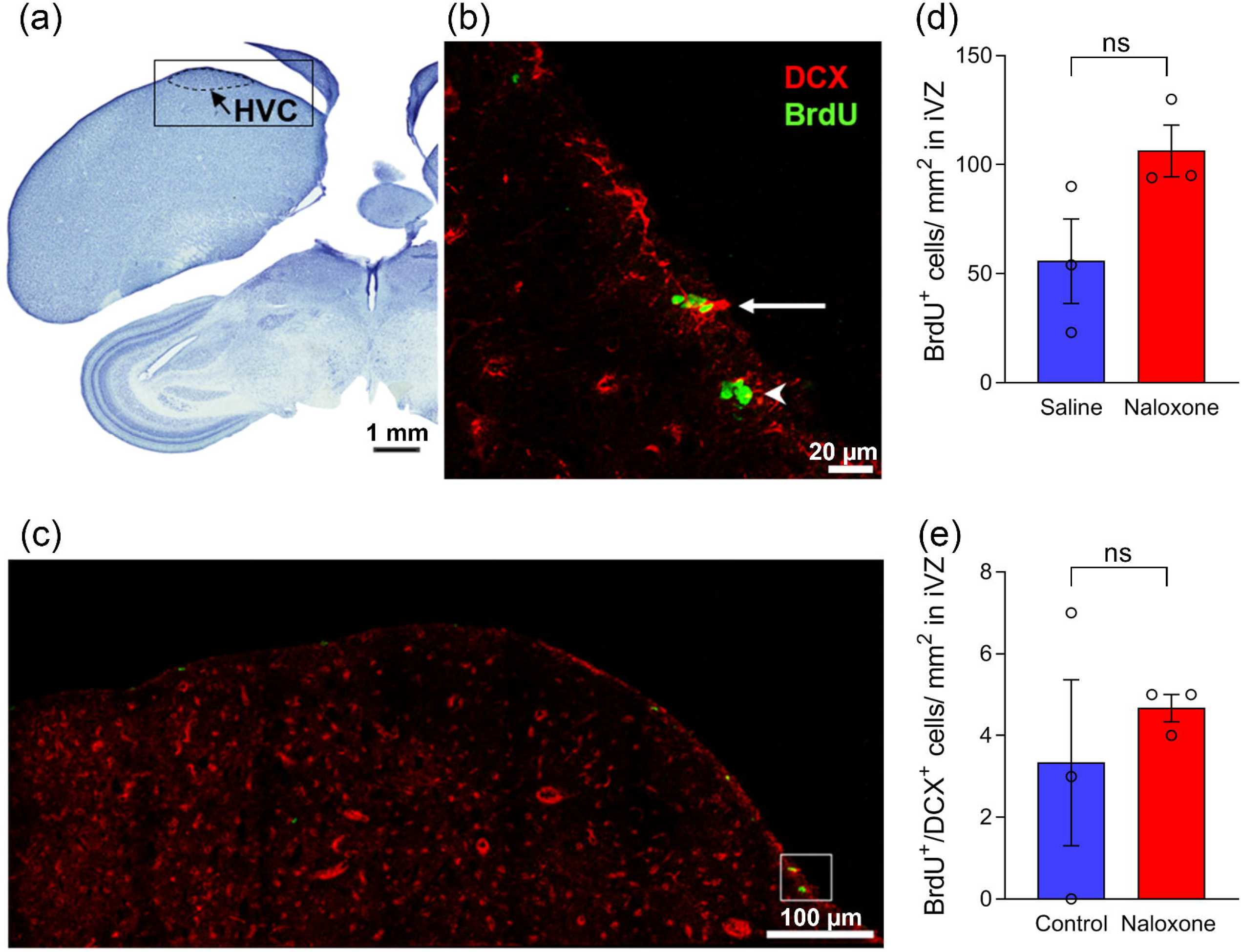
Blocking opioid receptors does not lead to changes in proliferation or neuronal differentiation in the VZ at the level of HVC. (**a**) A coronal section of the zebra finch brain stained with Nissl demonstrating the VZ dorsal to HVC (black rectangle); scale bar = 1 mm. **(b)** An image of the IVZ adjacent to HVC at low power; scale bar=100 µm and **(c)** the inset in (**b**) at a higher magnification (scale bar = 20µm) demonstrates dividing cells labeled with BrdU (*arrowhead*) and those co-labeled with BrdU and DCX (*arrow*). No significant increase was observed **(d)** in the number of BrdU+ cells/ mm^2^ (n = 3) or **(e)** co-labeled BrdU/DCX cells/mm^2^ (n = 4) in the IVZ dorsal to HVC of treated birds compared to that in controls. (Unpaired t-test, bars represent means ± SE, p<0.05)

## 5 Discussion and Conclusions

### 5.1 The Endogenous Opioid System modulates gene expression in the V-SVZ of Adult Male Zebra Finches

A whole transcriptome microarray on the adult male zebra finch V-SVZ revealed that systemic administration of the general opioid antagonist naloxone significantly upregulated the expression of 16 and downregulated the expression of 10 transcripts. Since the V-SVZ houses neural precursors, ependymal cells, neuroblasts, microglia and astrocytes [18], the differentially expressed transcripts observed in this study likely represent gene profiles of various cell populations present in the V-SVZ, which have responded to naloxone treatment. Furthermore, these results were validated by performing qRT-PCR analysis on selected transcripts, suggesting that the upregulation of these genes is indeed a consequence of opioid receptor blockade. Taken together, these findings suggest that the endogenous opioid system modulates gene expression the V-SVZ of adult male zebra finches.

### 5.2 Alterations in the Expression of Specific Genes in the V SVZ in Response to Naloxone Treatment

We found that the expression of the pre-miRNA *tgu-mir-124-201* transcript was significantly elevated in the V-SVZ of adult male zebra finches following naloxone treatment. The processing of precursor miRNAs into mature miRNAs involves the generation of duplexes containing two RNA strands, namely miR-124-3p and miR-124-5p [19], [20]. Subsequent qRT-PCR analysis of mature miRNA-124 in the neurogenic region after naloxone treatment confirmed the successful processing of *mir-124-201* into mature miRNA-124-3p and miR-124-5p. The observed upregulation of miR-124-3p (FC=1.6) but not of miR-124-5p (FC=1.1), in the naloxone-treated V-SVZ suggests that miR-124-3p is more stabilized and functionally relevant in this context.

MicroRNA-124 is the most abundantly expressed miRNA in the brain [21] and favours the differentiation of neural precursor cells into neurons [22]. A constant interplay between miRNAs and the endogenous opioid system must be in effect in order to maintain stable levels of neurogenesis in the brain [23], [24]. Our findings suggest that the levels of miR-124-3p, transiently upregulated during exposure to naloxone, favoured a shift towards the neuroblast stage from the neuronal precursor cells in the V-SVZ. These findings are supported by an earlier study which has demonstrated that miRNAs regulate mu-opioid receptors in rodents and humans [25]. Taken together, our findings underline the intimate involvement of mature miRNAs in modulating the effects of the endogenous opioid system on neurogenesis.

We also found that the transcript encoding Chromatin Accessibility Complex Subunit (CHRAC1) exhibited a 1.9-fold upregulation in the V-SVZ of naloxone-treated birds versus controls. CHRAC1 belongs to a group of DNA-binding histone-fold proteins [26], and plays a pivotal role in regulating the accessibility of DNA to the transcriptional machinery. Therefore, the higher expression of this gene suggests increased chromatin remodelling and transcriptional activation in the neurogenic zone of zebra finches after naloxone treatment.

Furthermore, there was a fold-change of 1.7 in the Adaptor Related Protein Complex 3 Subunit beta 2 (AP3B2) transcript in naloxone-treated V-SVZ samples compared to those of controls. Mutations of AP3B2 are known to be linked to microcephaly, developmental impediments, seizures, and intellectual disability [27], suggesting a potential role of AP3B2 in neuroblast migration and differentiation. Our results therefore suggest that blocking opioid receptors may affect AP3B2 expression in the V-SVZ, thereby modulating neurogenesis in adult male zebra finches.

Besides mir-124, we found that a transcript encoding TELO2 Interacting Protein 1 (TTI1) was upregulated by a fold change of 1.9 in the V-SVZ of naloxone-treated birds versus controls. TTI-1 interacts with mTOR (mechanistic Target of Rapamycin), a serine/threonine protein kinase of PIKK protein family and an essential regulator of translation, cell proliferation, and survival (reviewed in Ryskalin et al., 2017) An interplay between the µ-OR system and mTOR signalling pathway has been implicated in neuronal development, neuronal survival and synaptic plasticity [29]. Furthermore, the mTOR signalling cascade is activated in the auditory forebrain regions [caudomedial nidopallium (NCM) and caudomedial mesopallium (CMM) and the primary auditory forebrain (Field L)] of adult male zebra finches involved in song recognition learning [30], which are known to incorporate new neurons [31]. As a caveat, earlier studies demonstrated that general opioid antagonists led to the inactivation of the PI3K/AKT/mTOR pathway, thereby reducing cell proliferation in cancer cells [32] whereas opioid agonists like morphine promoted cell proliferation and migration in squamous cell carcinoma [33] via activating the PI3K/AKT/mTOR signalling pathway. However, in our experiment, four days of opioid antagonist treatment led to an increased expression of the TTI1 transcript, indicating the involvement of activated mTOR signalling cascade in the V-SVZ cells versus that in controls, which may reflect species differences.

### 5.3 Changes in Adult Neurogenesis as a result of altering Opioid Modulation in the Zebra Finch Brain

Our results demonstrated that naloxone administration induced an increase in cell proliferation (cf Khurshid et al., 2010) as well as an increase in the density of newly proliferated neuroblasts in the V-SVZ lining the lateral ventricles adjacent to the song control nuclei Area X and HVC, versus that in controls. However, our results indicated that the increase in density of newly proliferated cells and neuroblasts was significant only at the level of Area X, suggesting that altering opioid modulation not only affects cell proliferation but also promotes the generation of neuronal precursors. As a caveat, Chen et al. (2020) [34]demonstrated that primary neural stem cells (NSCs) in mouse embryos expressed low levels of opioid receptors during the first 53 days of differentiation. Furthermore, they found that naloxone was effective in promoting neurogenesis only during the initial stages (days 0-5) and inhibited neurogenesis during later stages (days 11-15) of differentiation and that naloxone promotes neurogenesis via pathways independent of opioid receptors. Although Khurshid et al. (2010) [10]had shown that opioid receptors are robustly expressed in the V-SVZ of adult male zebra finches, we cannot rule out the possibility of naloxone promoting neurogenesis by means of pathways other than by directly binding to opioid receptors. Additionally, there may be differences in the effects of naloxone on neurogenesis depending on the development age of NSCs from the V-SVZ as well as species differences.

Taken together, our results suggest that non-coding RNAs may be important for regulating neuronal proliferation and differentiation in the V-SVZ of zebra finches which leads to the addition of neurons in the song control region Area X. Future studies will focus on mechanistic details of how non-coding RNAs may contribute to song learning and vocalization in male songbirds.

## Acknowledgements

The authors are grateful to Dr Sarbani Samaddar, NBRC, Manesar, for her comments and suggestions on editing the initial drafts of the manuscript.

## References

[1] D. A. Lim and A. Alvarez-Buylla, “The Adult Ventricular–Subventricular Zone (V-SVZ) and Olfactory Bulb (OB) Neurogenesis,” Cold Spring Harb Perspect Biol, vol. 8, no. 5, pp. 18820–18821, May 2016, doi: 10.1101/CSHPERSPECT.A018820.

[2] L. C. Abbott and F. Nigussie, “Adult neurogenesis in the mammalian dentate gyrus,” Anat Histol Embryol, vol. 49, no. 1, pp. 3–16, Jan. 2020, doi: 10.1111/AHE.12496.

[3] J. M. García-Verdugo, S. Ferrón, N. Flames, L. Collado, E. Desfilis, and E. Font, “The proliferative ventricular zone in adult vertebrates: a comparative study using reptiles, birds, and mammals,” Brain Res Bull, vol. 57, no. 6, pp. 765–775, Apr. 2002, doi: 10.1016/S0361-9230(01)00769-9.

[4] M. F. Paredes, S. F. Sorrells, J. M. Garcia-Verdugo, and A. Alvarez-Buylla, “Brain size and limits to adult neurogenesis,” Journal of Comparative Neurology, vol. 524, no. 3, pp. 646–664, Feb. 2016, doi: 10.1002/CNE.23896.

[5] A. Alvarez-Buylla, M. Theelen, and F. Nottebohm, “Proliferation ‘hot spots’ in adult avian ventricular zone reveal radial cell division,” Neuron, vol. 5, no. 1, pp. 101–109, Jul. 1990, doi: 10.1016/0896-6273(90)90038-H.

[6] K. W. Nordeen and E. J. Nordeen, “Projection neurons within a vocal motor pathway are born during song learning in zebra finches,” Nature, vol. 334, no. 6178, pp. 149–151, Jul. 1988, doi: 10.1038/334149a0.

[7] F. Nottebohm, “Neuronal replacement in adulthood,” Ann N Y Acad Sci, vol. 457, no. 1, pp. 143–161, 1985, doi: 10.1111/J.1749-6632.1985.TB20803.X.

[8] N. Khurshid, V. Agarwal, and S. Iyengar, “Expression of μ- and δ-opioid receptors in song control regions of adult male zebra finches (Taenopygia guttata),” J Chem Neuroanat, vol. 37, no. 3, pp. 158–169, May 2009, doi: 10.1016/J.JCHEMNEU.2008.12.001.

[9] P. Parishar, N. Sehgal, and S. Iyengar, “The expression of delta opioid receptor mRNA in adult male zebra finches (Taenopygia guttata),” PLoS One, vol. 16, no. 8, p. e0256599, Aug. 2021, doi: 10.1371/journal.pone.0256599.

[10] N. Khurshid, L. S. Hameed, S. Mohanasundaram, and S. Iyengar, “Opioid modulation of cell proliferation in the ventricular zone of adult zebra finches (Taenopygia guttata),” The FASEB Journal, vol. 24, no. 10, pp. 3681–3695, Oct. 2010, doi: 10.1096/fj.09-146746.

[11] L.-C. Cheng, E. Pastrana, M. Tavazoie, and F. Doetsch, “miR-124 regulates adult neurogenesis in the subventricular zone stem cell niche,” Nat Neurosci, vol. 12, no. 4, pp. 399–408, Apr. 2009, doi: 10.1038/nn.2294.

[12] M. Akerblom et al., “MicroRNA-124 Is a Subventricular Zone Neuronal Fate Determinant,” Journal of Neuroscience, vol. 32, no. 26, pp. 8879–8889, Jun. 2012, doi: 10.1523/JNEUROSCI.0558-12.2012.

[13] X. Cao, S. L. Pfaff, and F. H. Gage, “A functional study of miR-124 in the developing neural tube,” Genes Dev, vol. 21, no. 5, pp. 531–536, Mar. 2007, doi: 10.1101/gad.1519207.

[14] V. DeWulf and S. W. Bottjer, “Neurogenesis within the juvenile zebra finch telencephalic ventricular zone: A map of proliferative activity,” J Comp Neurol, vol. 481, no. 1, pp. 70–83, Jan. 2005, doi: 10.1002/cne.20352.

[15] J. V. Aronowitz et al., “Unilateral vocal nerve resection alters neurogenesis in the avian song system in a region-specific manner,” PLoS One, vol. 16, no. 8, p. e0256709, Aug. 2021, doi: 10.1371/journal.pone.0256709.

[16] A. Lunde and J. C. Glover, “A versatile toolbox for semi-automatic cell-by-cell object-based colocalization analysis,” Sci Rep, vol. 10, no. 1, p. 19027, Nov. 2020, doi: 10.1038/s41598-020-75835-7.

[17] J. Polomova, K. Lukacova, B. Bilcik, and L. Kubikova, “Is neurogenesis in two songbird species related to their song sequence variability?,” Proceedings of the Royal Society B: Biological Sciences, vol. 286, no. 1895, 2019, doi: 10.1098/rspb.2018.2872.

[18] D. Mizrak et al., “Single-Cell Analysis of Regional Differences in Adult V-SVZ Neural Stem Cell Lineages,” Cell Rep, vol. 26, no. 2, pp. 394-406.e5, Jan. 2019, doi: 10.1016/j.celrep.2018.12.044.

[19] K. B. Choo, Y. L. Soon, P. N. N. Nguyen, M. S. Y. Hiew, and C.-J. Huang, “MicroRNA-5p and -3p co-expression and cross-targeting in colon cancer cells,” J Biomed Sci, vol. 21, no. 1, p. 95, Dec. 2014, doi: 10.1186/s12929-014-0095-x.

[20] Q. Li, S. Liu, J. Yan, M.-Z. Sun, and F. T. Greenaway, “The potential role of miR-124-3p in tumorigenesis and other related diseases,” Mol Biol Rep, vol. 48, no. 4, pp. 3579–3591, Apr. 2021, doi: 10.1007/s11033-021-06347-4.

[21] W.-H. Zhang, L. Jiang, M. Li, and J. Liu, “MicroRNA-124: an emerging therapeutic target in central nervous system disorders,” Exp Brain Res, vol. 241, no. 5, pp. 1215–1226, May 2023, doi: 10.1007/s00221-022-06524-2.

[22] N. A. Maiorano and A. Mallamaci, “The pro-differentiating role of miR-124: Indicating the road to become a neuron,” RNA Biol, vol. 7, no. 5, pp. 528–533, Sep. 2010, doi: 10.4161/rna.7.5.12262.

[23] H.-I. Im and P. J. Kenny, “MicroRNAs in neuronal function and dysfunction,” Trends Neurosci, vol. 35, no. 5, pp. 325–334, May 2012, doi: 10.1016/j.tins.2012.01.004.

[24] A. C. W. Smith and P. J. Kenny, “MicroRNAs regulate synaptic plasticity underlying drug addiction,” Genes Brain Behav, vol. 17, no. 3, Mar. 2018, doi: 10.1111/gbb.12424.

[25] S. L. Grimm et al., “MicroRNA–mRNA networks are dysregulated in opioid use disorder postmortem brain: Further evidence for opioid-induced neurovascular alterations,” Front Psychiatry, vol. 13, Jan. 2023, doi: 10.3389/fpsyt.2022.1025346.

[26] A. R. Mansisidor and V. I. Risca, “Chromatin accessibility: methods, mechanisms, and biological insights,” Nucleus, vol. 13, no. 1, pp. 236–276, Dec. 2022, doi: 10.1080/19491034.2022.2143106.

[27] M. Assoum et al., “Autosomal-Recessive Mutations in AP3B2, Adaptor-Related Protein Complex 3 Beta 2 Subunit, Cause an Early-Onset Epileptic Encephalopathy with Optic Atrophy,” The American Journal of Human Genetics, vol. 99, no. 6, pp. 1368–1376, Dec. 2016, doi: 10.1016/J.AJHG.2016.10.009.

[28] L. Ryskalin et al., “mTOR-Dependent Cell Proliferation in the Brain,” Biomed Res Int, vol. 2017, pp. 1–14, 2017, doi: 10.1155/2017/7082696.

[29] R. D. Polakiewicz, S. M. Schieferl, A. C. Gingras, N. Sonenberg, and M. J. Comb, “μ-Opioid Receptor Activates Signaling Pathways Implicated in Cell Survival and Translational Control,” Journal of Biological Chemistry, vol. 273, no. 36, pp. 23534–23541, Sep. 1998, doi: 10.1074/JBC.273.36.23534.

[30] S. Ahmadiantehrani, E. O. Gores, and S. E. London, “A complex mTOR response in habituation paradigms for a social signal in adult songbirds,” Learning & Memory, vol. 25, no. 6, pp. 273–282, Jun. 2018, doi: 10.1101/lm.046417.117.

[31] C. L. Pytte, S. George, S. Korman, E. David, D. Bogdan, and J. R. Kirn, “Adult Neurogenesis Is Associated with the Maintenance of a Stereotyped, Learned Motor Behavior,” The Journal of Neuroscience, vol. 32, no. 20, p. 7052, May 2012, doi: 10.1523/JNEUROSCI.5385-11.2012.

[32] N. Liu et al., “Low-dose naltrexone plays antineoplastic role in cervical cancer progression through suppressing PI3K/AKT/mTOR pathway,” Transl Oncol, vol. 14, no. 4, p. 101028, Apr. 2021, doi: 10.1016/j.tranon.2021.101028.

[33] L. Levi et al., “Effect of Opioid Receptor Activation and Blockage on the Progression and Response to Treatment of Head and Neck Squamous Cell Carcinoma,” J Clin Med, vol. 12, no. 4, p. 1277, Feb. 2023, doi: 10.3390/jcm12041277.

[34] J. Chen et al., “Naloxone regulates the differentiation of neural stem cells via a receptor-independent pathway,” The FASEB Journal, vol. 34, no. 4, pp. 5917–5930, Apr. 2020, doi: 10.1096/fj.201902873R.

