## Supplementary Tables 1-4 for "Opioid Modulation of Differential Gene Expression and Neuronal Differentiation in the Ventricular-Subventricular Zone of Adult Male Zebra Finches"

**Supplementary Table 1: Summary of differential expression analysis.**

| Conditions | Number of probes scanned | Number of filtered probes | Total no. of differentially expressed transcripts | Up-regulated Transcripts FC ≥ 1.5 | Down-regulated Transcripts FC ≥ 1.5 |
|---|---|---|---|---|---|
| Saline controls versus naloxone treated birds | 23,395 | 22,280 | 427 | 16 | 10 |

**Supplementary Table 2. Upregulated transcripts (p* < 0.05, |FC| ≥ 1.5) in the ventricular zone of adult male zebra finches administered with naloxone** (Highlighted genes have been validated by qRT-PCR)

| Ensembl Transcript ID | Gene Assignment | Fold Change | p-value |
|---|---|---|---|
| ENSTGUT00000015881 | Chromatin Accessibility Complex Subunit 1 (CHRAC1) | 1.9 | 0.038 |
| ENSTGUT00000014267 | TELO2 Interacting Protein 1 (TTI1) | 1.9 | 0.007 |
| ENSTGUT00000014673 | PHD finger protein 7-like | 1.8 | 0.035 |
| ENSTGUT00000018522 | Pre-MicroRNA 124-2(mir-124-201) | 1.7 | 0.013 |
| ENSTGUT00000014700 | Adaptor Related Protein Complex 3 Subunit beta 2 (AP3B2) | 1.7 | 0.037 |
| ENSTGUT00000010765 | NADH dehydrogenase [ubiquinone] 1 beta subcomplex subunit 3 (NDUFB3 ) | 1.6 | 0.041 |
| ENSTGUT00000001050 | Spliceosome-associated protein SPF 27 | 1.6 | 0.007 |
| ENSTGUT00000015827 | GRB2 related adaptor protein 2 | 1.6 | 0.002 |
| ENSTGUT00000013369 | Tumour necrosis factor alpha-inducible protein 2 (TNFAIP2) or Primary response gene B94 protein | 1.5 | 0.003 |

| | | | |
|---|---|---|---|
| ENSTGUT00000011073 | Uncharacterized protein C2orf66 Precursor | 1.5 | 0.003 |
| ENSTGUT00000000592 | Serine Peptidase Inhibitor, Kazal type 5 (SPINK5) | 1.5 | 0.002 |
| ENSTGUT00000015455 | Heme Binding Protein 1 (HEBP1) | 1.5 | 0.036 |
| ENSTGUT00000016722 | Novel Transcript | 1.5 | 0.024 |
| ENSTGUT00000009329 | Transmembrane Protein 180 | 1.5 | 0.017 |
| ENSTGUT00000014802 | Growth Hormone Releasing Hormone Receptor (GHRH) | 1.5 | 0.009 |
| ENSTGUT00000017025 | Contactin 6 (CNTN6) | 1.5 | 0.0002 |

**Supplementary Table 3: Downregulated transcripts ($P < 0.05$, $|FC| \geq 1.5$) in the ventricular zone of adult male zebra finches administered with naloxone** (Genes highlighted in bold been validated using qRT-PCR).

| Ensembl Transcript ID | Gene Assignment | Fold change | p-value |
|---|---|---|---|
| ENSTGUT00000018437 | uncharacterised novel transcript | 3.8 | 0.027 |
| ENSTGUT00000014358 | Induced myeloid leukemia cell differentiation protein MCL-1 | 1.8 | 0.009 |
| ENSTGUT00000012348 | uncharacterised novel transcript | 1.8 | 0.043 |
| ENSTGUT00000017353 | Related to Haloacid Dehalogenase like hydrolase domain containing 2 (HDHD2) | 1.7 | 0.017 |
| ENSTGUT00000018645 | uncharacterized LOC116808170 non-coding RNA | 1.6 | 0.032 |
| ENSTGUT00000018842 | C19orf12 homolog | 1.6 | 0.048 |
| ENSTGUT00000019351 | SNORA74 // Small nucleolar RNA | 1.6 | 0.009 |

| | | | |
|---|---|---|---|
| ENSTGUT00000019188 | chromosome 22 open reading frame 41 | 1.5 | 0.003 |
| ENSTGUT00000004096 | related to RNA helicase Mov10l1 | 1.5 | 0.035 |
| ENSTGUT00000019348 | Small nucleolar RNA SNORD26 | 1.5 | 0.048 |

**Supplementary Table 4: Sequences of primers used in the qRT-PCR analysis.**

| Gene | | 5' to 3' |
|---|---|---|
| TNF alpha induced protein 2 (TNFAIP2) | Forward | CTCCCAGGCAAAGAAAAATG |
| | Reverse | TAAAAGTTGTGCACCCACGA |
| Chromatin accessibility complex subunit 1 (CHRAC1) | Forward | AGCGAGATGGAGAAATGAGG |
| | Reverse | AACAGCAGCACCAACACCTT |
| TELO2 interacting protein 1 (TTI1) | Forward | GAGGCCTGTGGTTACGACTC |
| | Reverse | AATTCAGGGAAATCCCGTTC |
| Adaptor related protein complex 3 subunit beta 2 (AP3B2) | Forward | GGGAGCCTGGTACTCATCAC |
| | Reverse | TGACCATCTTCTCGCTGTTG |
| Induced myeloid leukemia cell differentiation protein MCL-1 | Forward | CAG CCC AGC GTT AAG AAG |
| | Reverse | CTT CTC CAT CAC CGC ATC |
